# Time to recover from daily caffeine intake

**DOI:** 10.1101/2021.07.01.450708

**Authors:** Yu-Shiuan Lin, Janine Weibel, Hans-Peter Landolt, Francesco Santini, Corrado Garbazza, Joshua Kistler, Sophia Rehm, Katharina Rentsch, Stefan Borgwardt, Christian Cajochen, Carolin Reichert

## Abstract

Caffeine elicits widespread effects in the central nervous system and is the most frequently consumed psychostimulant worldwide. First evidence indicates that, during daily intake, the elimination of caffeine may slow down and the primary metabolite, paraxanthine, may accumulate. The neural impact of such adaptions is virtually unexplored. In this report, we leveraged the data of a laboratory study with N= 20 participants and three within-subject conditions: caffeine (150 mg caffeine x 3/day x 10 days), placebo (150 mg mannitol x 3/day x 10 days), and withdrawal (caffeine x 9 days, afterwards placebo x 1 day). Using liquid chromatography–mass spectrometry coupled with tandem mass spectrometry, we determined the course of salivary caffeine and paraxanthine, measured regularly at day 10. We assessed grey matter (GM) intensity and cerebral blood flow (CBF) in the withdrawal condition as compared to their changes in caffeine in our previous report. The results indicate high remaining levels of paraxanthine and of caffeine carried overnight during daily intake, and the levels of paraxanthine remained higher than in placebo during withdrawal. After 36 h of withdrawal, the previously reported caffeine-induced GM reduction was partially mitigated, while CBF was elevated compared to placebo. Our findings unveil that conventional daily caffeine intake does not provide sufficient time to clear up psychoactive compounds and restore cerebral responses, even after 36 hours of abstinence. They also suggest investigating consequences of a paraxanthine accumulation during daily caffeine intake.

## Introduction

Caffeine is the most frequently consumed psychostimulant worldwide. After an acute caffeine administration, 99% of the administered caffeine dose can be absorbed in ∼45 min (1), which brings the peak plasma concentration of caffeine at 1 – 2 h after the intake. The average half-life of caffeine after an acute consumption is 2.5 – 5 h (2), while it can be modulated by the ingested dosages, smoking, genetic variance, health status, oral contraceptives and pregnancy, and various other factors (3-6). After absorption, around 80% of caffeine is metabolized into paraxanthine through majorly the cytochrome P450 1A2 enzyme (CYP1A2) in the liver. Following the peak concentration after a single administration of 5–8 mg/kg caffeine, plasma concentrations of caffeine decline rapidly and are surpassed by the paraxanthine levels at around 8 – 10 h after the intake (7). Paraxanthine has as high potency at antagonizing adenosine receptors as caffeine (8) and exerts several similar effects as caffeine, such as wake-promotion (9), psychostimulation (10), elevating blood pressure, and release of epinephrine (11). Yet, kinetics and concentrations of paraxanthine have rarely been considered when studying physiological and cognitive outcomes of caffeine intake in humans.

Despite the common pattern of regular intake, studies on the kinetics of caffeine and paraxanthine in humans after daily caffeine intake are surprisingly scarce. In rodents, applying caffeine for 10 days in a 3-h interval leads to a smaller elimination rate constant (K_el_) of plasma caffeine concentration after 10 days of caffeine intake compared to acute administration (12). Furthermore, compared to a linear dose response in a food-limited state, daily caffeine intake in an *ad lib* dietary state leads to dose-disproportional responses (i.e. the higher the dose the larger increase) in peak levels of caffeine and paraxanthine and the 24-h area under the curve (AUC_0-24_) (12). In humans, only one study compared the metabolism of caffeine in nine adults among baseline (0 mg/kg), low-dose (0.7mg/kg), and high-dose (2 mg/kg) treatments applied every 2 h, 6 times a day for 5 days (4). The authors reported a dose-dependent deceleration of metabolism of intravenous isotope-labeled (2-’3C, 1,3-’5N2) caffeine after three days of treatment (i.e., the higher the dose, the slower the metabolism), and a dose-disproportional elevation in the AUC_0-24_ of paraxanthine. This study, however, included a rather small sample size with an unusually high dosage, given that the volunteer with highest weight of 99 kg could receive up to 1188 mg/day in the high-dose condition. It remains unclear whether the metabolism during a conventional pattern of daily caffeine intake over a longer time will adapt similarly as in this study. We consider a treatment with a unified dose in the morning, noon, and afternoon hours as more generalizable (13-15).

Furthermore, although these findings consistently suggest adapted metabolism of caffeine and paraxanthine over the course of daily caffeine intake, no physiological outcomes were available in these studies. An increase of psychoactive compounds may also multiply the impacts in brains, as *ex vivo* evidence indicates that elevating caffeine concentrations can lead to an increase in the brain-to-plasma ratio of caffeine (16). Previously, we observed a decreased grey matter (GM) in hippocampus and reduced cerebral blood flow (CBF) during daily caffeine intake, in which the larger reductions of both properties were associated with a higher accumulation of caffeine + paraxanthine. We postulated that these cerebral responses may be due to an incomplete elimination and an accumulation of the psychoactive compounds. It is still unclear, however, how fast these brain responses can be restored during abstinence, when caffeine and paraxanthine can be completely eliminated.

In addition to the daily pattern of intake, genetic variations also determine individual metabolic process and in turn modulate the development of habitual patterns of caffeine intake (5, 17, 18). In particular, the variants in CYP1A2, AHR, and CYP2A6, which are associated with lower habitual intake (19, 20), are also associated with slower metabolism of caffeine (CYP1A2 and AHR) and paraxanthine [CYP2A6; (19)]. Thus, the adaptions in the metabolism of caffeine and paraxanthine during daily caffeine intake and its physiological effects can be also modulated by the genetic traits reflected by the levels of habitual intake.

For the present report, we leveraged salivary samples of a previous study with three within-subject conditions (21): caffeine (150 mg caffeine x 3/day x 10 days), placebo (150 mg mannitol x 3/day x 10 days), and withdrawal (caffeine x 9 days + placebo x 1 day). We report a 43-h profile of caffeine and paraxanthine levels, the kinetics of each compound and their association with habitual caffeine intake. Moreover, we examined the brain recovery after 36 h of withdrawal when the levels of caffeine were expected to be cleared. The protocol simulated a conventional pattern of a double espresso at breakfast, lunch, and afternoon-teatime (in 3.25- and 4-h interval, respectively).

## Methods

### Study protocol, participants, and environmental control

In a double-blind, randomized, placebo-controlled study, we included twenty clinically healthy male non-smokers, who were between 18 and 35 years old with a body mass index between 18 and 26and a habitual caffeine intake between 300 and 600 mg/day. The amount of habitual caffeine intake was assessed by a self-report questionnaire adapted from Bühler, Lachenmeier (22) and Snel and Lorist (23).

Each volunteer completed three conditions (orders of condition see (24)): A placebo condition (150 mg mannitol x 3/ day x 10 days), a daily caffeine condition (150 mg caffeine x 3/ day x 10 days), and a caffeine withdrawal condition (caffeine until the first administration of the 9^th^ day, then switch to placebo until the end of 10^th^ day). The outcomes were assessed on the 10^th^ day of each condition. **Figure 1** presents the study protocol, including the timing of the treatments, the outcome measurements, and the corresponding hours of withdrawal in each condition. We scheduled the magnetic resonance imaging (MRI) scans in all three conditions for all participants at 12 h after awakening, in order to control for the confounding effects of different durations of wakefulness (25) and different levels of sleepiness (26) on cerebral variables.

**Figure 1.**
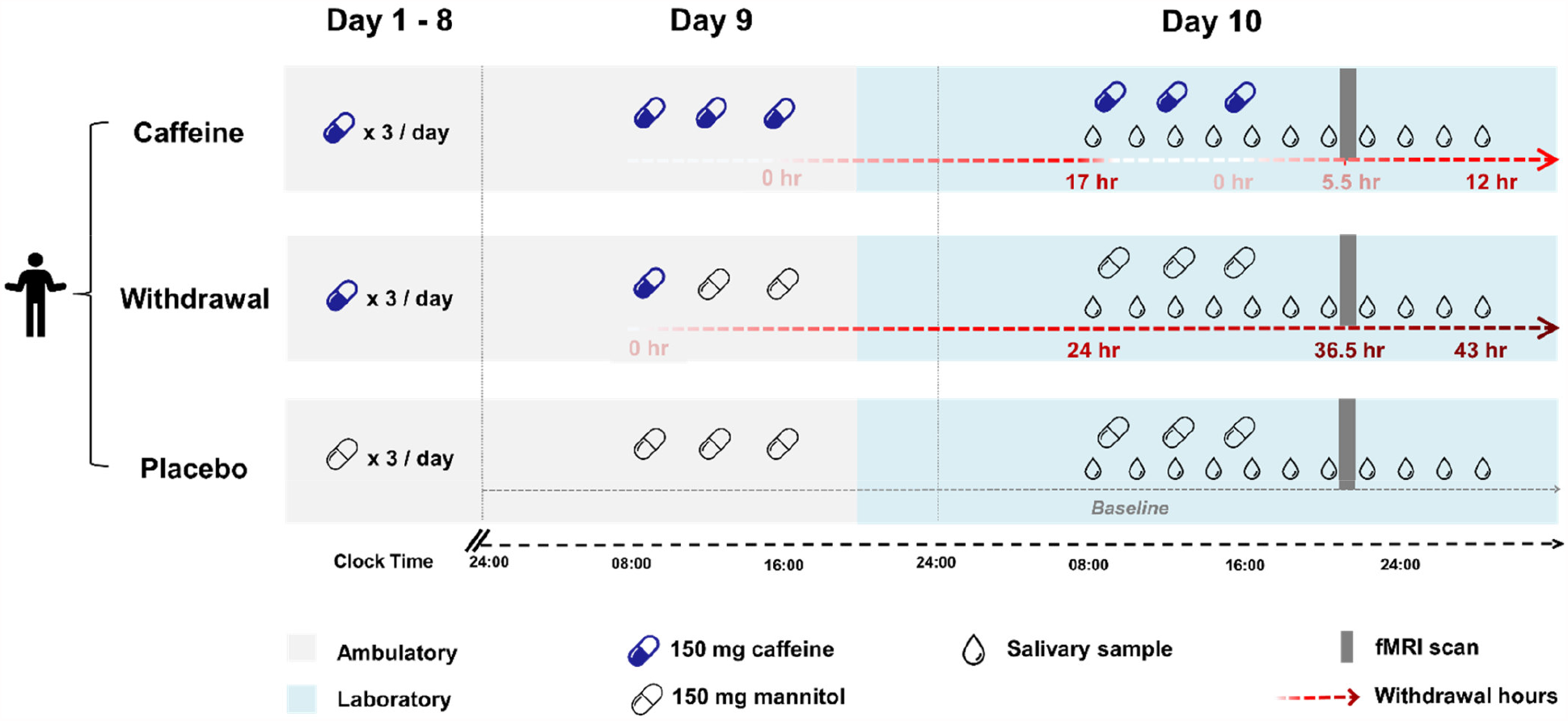
Overview of study design. Each condition consisted of 9 ambulatory days (grey shading) and a subsequent 21-h laboratory stay starting in the evening of day 9 (blue shading). Timing of caffeine or placebo intake is indicated by capsules (blue: caffeine, transparent: placebo), timing of saliva collection to measure caffeine and paraxanthine levels is indicated by drops. ***Top:*** Caffeine condition: caffeine capsules (150 mg caffeine) x 3 times/ day (at 45 min, 4 h, and 8 h after awakening) were administered throughout 10 days. Saliva was collected repeatedly during the laboratory stay from right before the first capsule until 12 h after the latest caffeine intake. The MRI scan (grey bar) took place at 5.5 h after the latest caffeine intake. ***Middle:*** Withdrawal condition: caffeine capsules (150 mg caffeine) x 3 times/ day were administered for 8 days, followed by a switch to placebo capsules on day 9. Saliva was collected in between 24 and 43 h after the latest caffeine intake. The MRI scan was scheduled to 36.5 h after the latest caffeine intake, a time window of strong withdrawal symptoms (28). ***Bottom:*** Placebo condition: placebo capsules (150 mg mannitol) x 3 times/ day were administered throughout the ambulatory and laboratory days. Saliva was collected as baseline through the entire laboratory phase at the corresponding time points as the other two conditions. The MRI scan was scheduled at the same time of day as in the other two conditions.

Through the entire 10 days, participants complied with a regular sleep-wake cycle (8 h ± 30 min nighttime sleep, no naps allowed) according to their habitual bedtime, which was monitored and recorded by actimetry and self-reported sleep diaries. Participants abstained from any other caffeine-containing diets, including coffee, tea, energy drink, soda, and chocolate, etc., and the compliance was monitored by the caffeine levels in the evenings. The data of one participant’s caffeine condition was excluded due to incompliance with the treatment. The laboratory environment from the 9^th^ day evening through the 10^th^ day was strictly controlled: dim light, half-supine posture (∼ 45°), and regular dietary and lavatory times. Participants were allowed to sleep for 8 hours in the night between day 9 and day 10. Digital device without time clues and internet was permitted. Social interaction was restricted to the group of experimenters.

### Data acquisition and analyses

#### 1. Caffeine and paraxanthine

In each condition, we analyzed caffeine and paraxanthine levels in 11 saliva samples, collected in 105 – 120 min intervals. Salivary caffeine and its main metabolites (paraxanthine, theobromine, and theophylline) were analyzed by High Performance Liquid Chromatography (HPLC) coupled to tandem mass spectrometry at the Laboratory Medicine, University Hospital Basel. Values below the detection threshold of 20 ng were treated as 20 in the analyses to eliminate the nuisance from the stochastic variance.

We used the samples in caffeine condition to calculate the kinetics of caffeine and paraxanthine by peak level (Cmax), peak time (Tmax), elimination rates (K_el_), and half-lives. In addition, we characterize the accumulation of caffeine and paraxanthine by the AUC in caffeine (AUCc) and in the withdrawal (AUCw) condition. Furthermore, we examined the overnight residuals of each compound in the caffeine condition in the morning, right before the first treatment on Day 10. This level indicated the progress of the elimination of caffeine and paraxanthine during the repetition of daily intake. *Cmax* was defined by the maximal level after the last caffeine administration, and the latency to Cmax (h after the last intake) was used *Tmax*. Due to the sampling frequency, the earliest sample collected after the last administration was roughly by 105 min, which limited the quantification of the variances of Cmax and Tmax below this threshold. A *half-life* was defined by the time from the peak to the time when the concentration approximates 50% of the maximal level (C_50_). *AUC of caffeine and of paraxanthine* were calculated with the trapezoidal rule over the eleven samples separately per caffeine and withdrawal condition.

#### 2. GM and CBF

T1-weighted structural data were obtained with a Magnetization-Prepared Rapid Gradient Echo (MP-RAGE) sequence (1x1x1mm^3^, TR=2000ms, TE=3.37ms, FA=8°) on a 3T Siemens scanner (MAGNETOM Prisma; Siemens Healthineers, Erlangen, Germany). Cerebral blood flow (CBF) was measured by 2D Echo-Planar Imaging (EPI) pulsed arterial spin labeling (ASL) sequence (4x4x4mm^3^, TR=3000ms, TE=12ms, FA=90). For the detailed pipeline of preprocessing and the whole-brain analyses (WBAs), please find the methods in (21). The current analysis adopted a region-of-interest (ROI) approach based on the previously reported caffeine-induced results in a whole-brain analysis (21). The pre-defined regions are the right hippocampus for GM, and precuneus, thalamus, and basal ganglia for CBF. We extracted the mean intensity of GM (indicating both volume and density of GM) and the mean quantity of CBF in the respective ROIs in all three conditions and examined the recovery status in the withdrawal condition compared to placebo and caffeine.

#### 3. Statistical analysis

All the statistics were conducted on R (R core team, Vienna, Austria). To analyze the interaction effect between condition x time in caffeine and paraxanthine as well as the condition effects in regional intensity of GM and CBF, we first estimated the coefficients with a generalized linear mixed model by the *afex* package (27), followed by ANOVA to obtain the statistical parameters of variance analysis (F, T, and p values). Post-hoc pairwise analysis was performed by using Tukey’s multiple comparison test. The inter-individual analyses on associations between doses (including habitual caffeine intake and treatment dose) and half-lives of caffeine and paraxanthine were performed by generalized linear regressions.

## Results

### 43-h profiles and pharmacokinetics

Twenty volunteers (age: 26.4 ± 4.0 years; BMI: 22.7 ± 1.4 kg/m^2^; weights: 76.2 ± 8.7 kg; relative dosage of laboratory treatment: 6.0 ± 0.6 mg/kg/day; and self-reported habitual caffeine intake: 6.8 ± 2.3 mg/kg/day) completed the study. We present the 43-h profile of caffeine and paraxanthine in **Figure 2**, and the exhaustive data of all samples in three conditions are included in the supplement (**Table S1**). The significant main effect of condition (F_(2, 602)_ = 618.0, p < .001) on salivary levels of caffeine indicated overall higher levels in the caffeine condition (t = 30.5, p < .001) compared to placebo. The overall levels of caffeine in the withdrawal condition were significantly lower compared to caffeine condition (t--28.7, p < .001) but did not significantly differ from placebo (t = 1.7, p = .190). Indicated by the condition x sample interaction (F_(10, 602)_ = 9.3, p < .001), the overnight residual level of caffeine already *before* the first laboratory intake in the caffeine condition was significantly higher than in the placebo condition (see 17h after the latest intake in **Figure 2**, t = 3.6, p < .001). Caffeine levels in caffeine condition remained higher than in placebo condition until 12 h after the latest intake (t = 5.3, p < .001).

**Figure 2.**
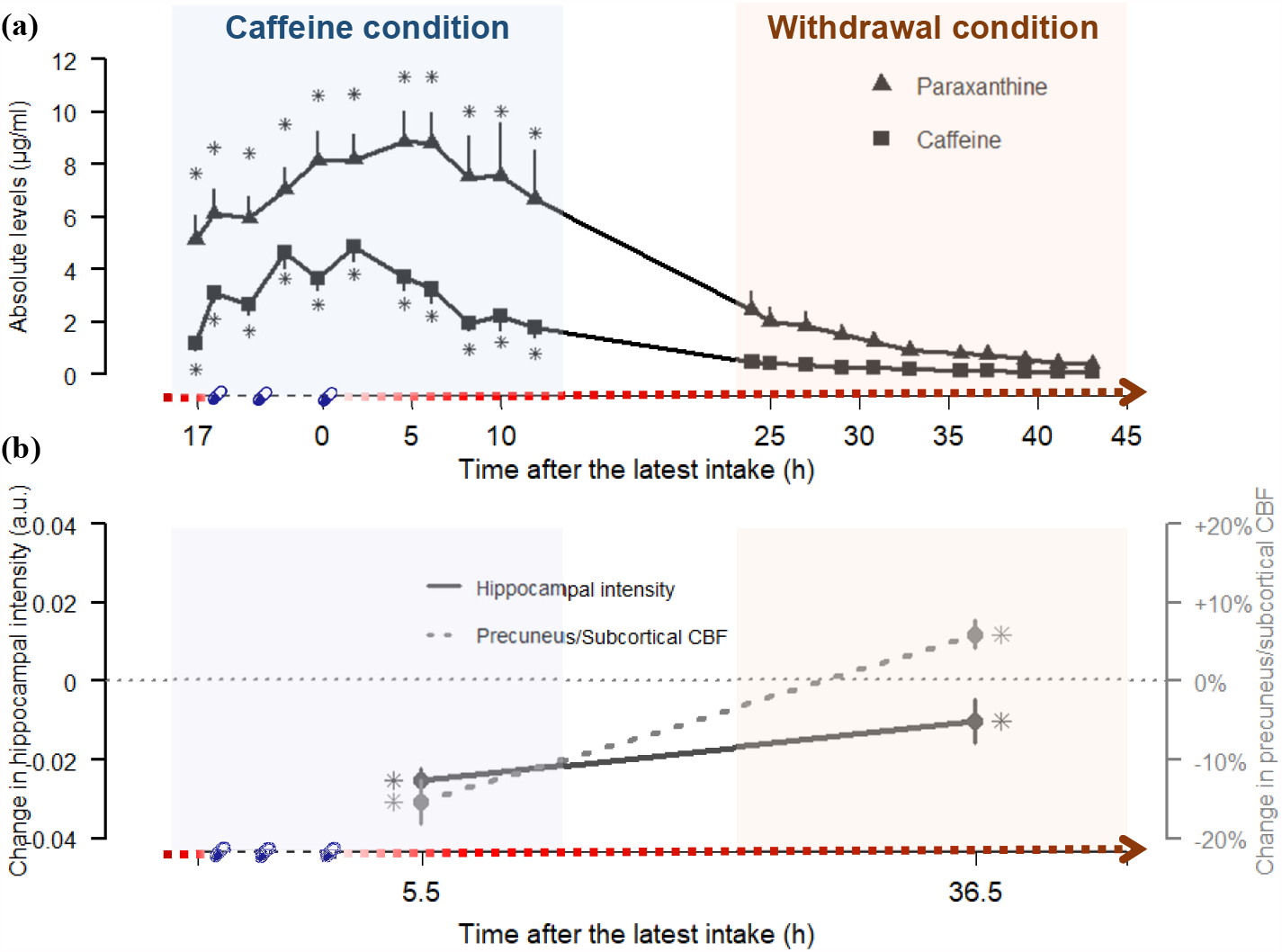
Overview of the paraxanthine and caffeine values at the respective time points and the dynamics of cerebral responses. Panel **(a)** juxtaposes the profiles of caffeine (squares) and paraxanthine (triangle) in caffeine (blue shading, left panel) and withdrawal conditions (red shading, right panel) against the time after the latest dose of caffeine on the x-axis. Echoing **Fig 1**, the gradient red arrow parallel to the x-axis schematically indicates the intensity of the withdrawal. The times of treatments are denoted by blue capsules. Detailed statistics are addressed in the result section. In brief, the main effects of conditions indicated that paraxanthine levels remained elevated throughout the caffeine and withdrawal conditions compared to placebo, while caffeine levels were elevated in caffeine condition but did not significantly differ from placebo in withdrawal. Asterisks (*) indicated the time points exhibiting significant elevated levels compared to placebo by a post-hoc analysis on the significant Condition x Sample interaction. Panel **(b)** illustrates the magnitudes of the changes in hippocampal GM intensity (straight line) and the precuneus + subcortical CBF quantity (dashed line)during daily caffeine intake and after caffeine withdrawal, relative to placebo (dotted horizontal line). The asterisks indicated significant differences compared to placebo. Detailed statistics are addressed in the result section.

Regarding the salivary levels of paraxanthine, we also observed a significant main effect of condition (F_(2, 602)_ = 400.4, p < .001). A post-hoc analysis indicated that the paraxanthine levels were significantly higher in the caffeine (t = 25, p < .001) than in the placebo condition, while in withdrawal, the levels were massively reduced (t = -21.7, p < .001, compared to caffeine) but still significantly higher than in placebo condition (t = 3.8, p < .001). Furthermore, indicated by the condition x sample interaction (F_(10, 602)_ = 2.0, p = .007), the overnight residual level of paraxanthine before the first laboratory intake in the caffeine condition was significantly higher than in the placebo condition (t = 5.4, p < .001). The paraxanthine levels remained higher than in the placebo condition after 12 h after last intake (t_C12_ = 6.9, p_C12_ <.001).

In **Table 1**, we present the pharmacokinetics of caffeine and paraxanthine, including peak times, peak levels, and half-lives after the laboratory caffeine administration, following the prior ambulatory 9-day consumption. Paraxanthine peaked 3.2 h later than caffeine (t = 3.8, p = .002) and had a 3.5 h longer half-life than caffeine (t = 2.7, p = 0.15). Furthermore, after correcting for the false discovery (reported in q values), the regression coefficients (**Figure 3**) indicated that higher habitual intake was significantly associated with longer half-life of caffeine (β = -0.11, p = .002, q = .006) and of paraxanthine (β = -0.14, p = .04, q = .04), as well as *at trend* associated with the larger disproportionality between paraxanthine and caffeine (AUCc of PX/AUCc of CA: β = 0.05, p = .036, q = .054).

**Table 1.**
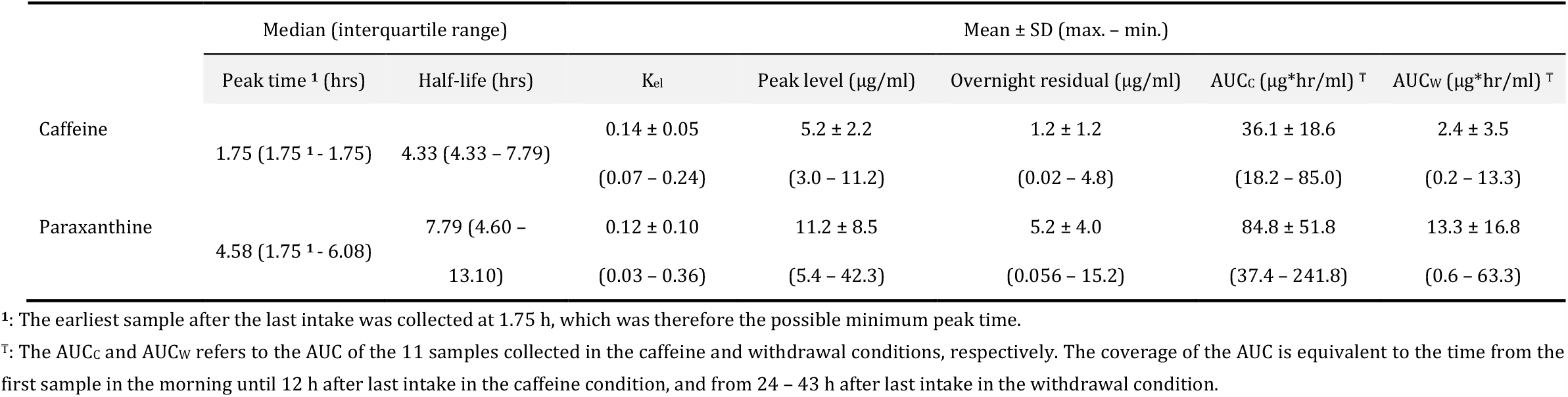
Peak level, peak time, half-life, and morning residuals of caffeine and of paraxanthine during the caffeine condition.

**Figure 3.**
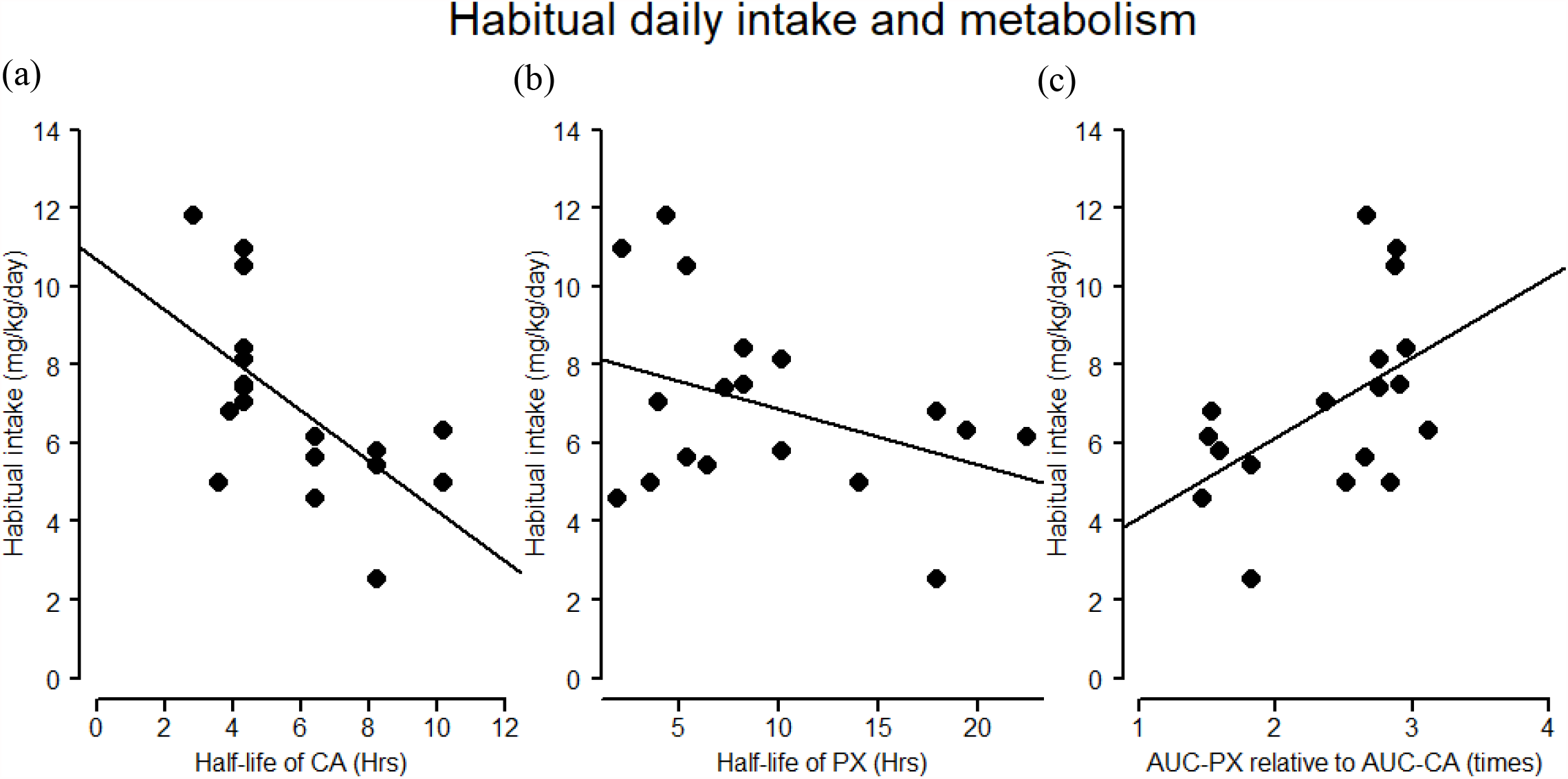
Associations of habitual daily intake and half-lives of caffeine and paraxanthine. (a) and (b) depict the negative association between habitual self-reported caffeine intake (mg/kg/day) and the half-life of caffeine and paraxanthine in the caffeine condition, respectively. (c) illustrates the association between habitual caffeine intake (mg/kg/day) and the ratio of AUC of paraxanthine (AUC-PX) to AUC of CA (AUC-CA). In (c), a high ratio of AUC-PX to AUC-CA indicates a larger disproportioanl accummulation between paraxanthine and caffeine.

### Brain recovery at 36 h of withdrawal

As illustrated by **Figure 2**, the reduced hippocampal GM during daily caffeine intake reported earlier elsewhere (21) remained significantly lower after 36.5 h of caffeine withdrawal compared to the placebo condition (t_W-P_ = -2.1, p_W-P_ = 0.039). The magnitude, however, was reduced in withdrawal compared to the caffeine condition (t_W-C_ = 3.2, p_W-C_= 0.003). On the other hand, CBF in precuneus, thalamus, and basal ganglia, which was reduced during daily caffeine intake (21), was elevated at 36.5-h in the withdrawal compared to the placebo condition (t_W-P_ = 2.1, p_W-P_ = 0.039).

## Discussion

The present study provides the first estimates for the kinetics of caffeine and paraxanthine and the responses of GM and CBF during and after a typical daily caffeine intake pattern [i.e., intake in the morning, at noon, and in the afternoon (13-15)]. We observed elevated residuals of paraxanthine and of caffeine carried overnight during the daily intake, suggesting an accumulation over repeated daily caffeine consumption. In line with earlier studies, the *at trend* association between the habitual caffeine intake and the proportion of AUC-PX to AUC-CA indicated people with higher habitual intake may tend to have a larger accumulation of paraxanthine during caffeine metabolism than lower habitual consumers. Additionally, our data allowed for the first time characterizing a recovery from daily intake (i.e. withdrawal) during 24-43 hours after the latest caffeine intake, where the overall paraxanthine levels remained elevated. Lastly, the caffeine-induced reduction in hippocampal GM intensity was only partially recovered by 36 h of withdrawal. In contrast, CBF was significantly elevated compared to placebo. Taken together, our data suggest that conventional daily caffeine intake does not provide sufficient time for an elimination of caffeine and paraxanthine. Furthermore, even though the salivary caffeine nearly reaches a clear state, the caffeine-associated brain responses will require longer time to be fully restored. Finally, the accumulation of paraxanthine entails a critical role in the effects of caffeine consumption in chronic intakes. Given its high potency at the cerebral adenosine receptors (8), the potential effects of prolonged exposure to this xanthine and its clinical application should be further inspected in the future.

### Daily moderate caffeine and accumulation of paraxanthine

Daily repeated intake of caffeine is a very common phenomenon and occurs in adults more frequently as compared to an acute irregular consumption of the psychostimulant after a certain phase of abstinence [patterns of regular caffeine intake see (14)]. Beside the present data, however, the evidence on the course of human caffeine metabolism under conditions of daily intake is scarce. Compared with earlier studies, our data however showed a more similar profile of caffeine metabolism to the evidence derived from acute (7, 29-32), but not repeated, intake (4).

With a unified dose of 150 mg x 3/day, our study had an average relative dose of 2 mg/kg x 3 times/day in a 4-h interval. As a result, the median peak time of caffeine (<105 min) in this study was similar to peak times observed both after acute (7, 29-32) and daily administration (4). The peak time of paraxanthine (4.6 h), however, was 3 h longer than the earlier evidence derived from a similar *total daily* dose (0.7 mg/kg x 6/day) while rather similar to the peak time from a similar single-dose intake (2 mg/kg x 6/day) in an earlier study (4). It is possible that, rather than a *total daily* dose level, the peak time of paraxanthine during repeated intake may depend more on the *single* dose level as well as the subsequent transformation of caffeine to paraxanthine.

On the other hand, the observed half-life of caffeine (4.3 h) is also similar to those after an acute administration [ranged between 2 - 6 h (7, 29-32)]. Compared to the aforementioned study using repeated daily doses, however, the half-lives of both caffeine and paraxanthine (7.8 h) in our data were much shorter, irrespective of different doses and timing of administration (4). The shorter half-life could be attributed to a longer interval (4 h) between the repetition of doses compared to 2 h in Denaro, Brown (4), as the longer interval in our study was nearly the half-life time of caffeine, which may have reduced the iteration of the caffeine concentration over repeated intake. Moreover, as our study only included male participants, a slower *average* metabolism of caffeine and paraxanthine in the earlier study could also in part be traced back to the influence of estrogen levels or the use of oral contraceptives in female participants (5). In addition, a faster metabolism in our study can also reflect genetic traits biased by our inclusion criterion on a habitual intake level of 300 – 600 mg/day. Supported by our observation of a negative association between habitual intake and half-lives of caffeine and paraxanthine, our selection of participants may also contribute to the discrepancy between the average caffeine metabolisms in our and the earlier study (4).

Furthermore, the concentration of paraxanthine in our study exceeded caffeine levels throughout the entire course, which was not seen in other human studies with a daily caffeine treatment (4, 33). The paraxanthine concentration in Baur, Lange (33), however, did show an ascending trend over 4 days of 300 mg/day intake. Suggested by the *at trend* association of a higher habitual intake to a more severe disproportionality between the accumulation of paraxanthine and of caffeine, this phenomenon in our data may result from a personal trait that determines a faster transformation of caffeine to paraxanthine without commensurate speed of metabolism of paraxanthine. In addition, we could not exclude the possibility of an existing accumulation from the participants’ habitual intake prior to the study, as this study did not implement a prior washout period before the start of ambulatory treatments. Nevertheless, the constant exceedance of paraxanthine levels over caffeine reveals an explicit risk of habitual or daily caffeine intake for a massive accumulation of paraxanthine. In fact, a dose-disproportional elevation in paraxanthine is not limited to daily intake. A similar phenomenon was also found in acute caffeine administration, where increasing caffeine concentration could multiply the increase of paraxanthine levels (34).

Overall, we observed an accumulation of paraxanthine present during a pattern of caffeine intake, which imitated a realistic dose and timing of consumption. This accumulation was further intensified in people with a higher dose of habitual caffeine consumption. Paraxanthine is considered to possess a lower toxicity and anxiogenesis than caffeine and other methylxanthines (9, 35, 36), while it can still exert similar effects as caffeine in wake-promotion (9), psychostimulation (10), elevating blood pressure, and release of epinephrine (11). Since paraxanthine levels are prone to be mounted over daily intake, understanding the cognitive and physiological outcomes in prolonged exposure to paraxanthine will clarify the long-term benefits and harms of daily caffeine intake.

### Do we sufficiently recover during daily caffeine intake?

Withdrawal responses commonly occur after discontinuing regular caffeine intake. In the current analyses, we corroborated the withdrawal-induced vasodilation by the elevated cerebral blood flow in the withdrawal condition (37-42). Beside the neurovascular effect, the cognitive responses in the same volunteers reported elsewhere, including reduced vigilance, increased sleepiness, and enhanced sleep depth (24, 43), confirmed that the participants were experiencing a solid withdrawal state.

Withdrawal responses can manifest as a restoration of adapted neural functioning when caffeine intake is ceased. The molecular mechanism of such adaptions include a mounting concentration of extracellular endogenous adenosine (44) and upregulated adenosine receptors (45-48). Both mechanisms can lead to a surge of cognitive and physiological reactions through the strengthened adenosine binding. These reactions include fatigue, reduced concentration, mood disturbance, headache, as well as increased vasodilation as addressed (28, 49, 50). Intense withdrawal symptoms are usually perceived around 20 and 50 h after the last regular intake and, in extreme cases, can last maximal 9 days (28). The earliest observed neurovascular responses to caffeine withdrawal in existing evidence was at 21 h (40).

While combating withdrawal symptoms are often the reason to consolidate the daily repeated consumption of caffeine (51), the typical daily repeated caffeine intake, however, is unlikely to provide enough time for a full “withdrawal-driven restoration”. The first evidence comes from our observation in the overnight residuals of caffeine and paraxanthine levels, which were measured at 17 h after the last intake. Furthermore, in the withdrawal condition (i.e. 24 h - 43 h of withdrawal) when the caffeine levels were nearly cleared, the paraxanthine levels remained elevated. In other words, a repetition of intake shorter than this time window is most likely to be insufficient for full elimination of both caffeine and paraxanthine. The prolonged presence of high levels of such adenosine antagonists may impede the neural homeostasis dependent on the activations of adenosine receptors. In neurogenesis of adult rodents, the presence of a similar dose of caffeine during sleep has been shown to suppress the cell proliferation in hippocampus [(52); Dose conversion: 10 mg/kg in rodents were estimated to be equivalent to 250 mg/ 70 kg in humans (51)]. The concurrence of a reduced hippocampal GM intensity and an incomplete caffeine elimination, together with a GM recovery in caffeine withdrawal, underscore the importance of a sufficient withdrawal period. The overcompensation of the CBF response in the withdrawal condition, again, points to a longer time – perhaps some consecutive days – for full recovery. Future studies should confirm this postulation with a design which allows full elimination caffeine and paraxanthine as well as an observation of the response in cerebral morphology.

## Limitations

This study bears a few limitations which should be carefully taken into account. First, as discussed earlier, the current analysis leveraged the data collected in a study without a washout period prior to the start of ambulatory treatment. This may limit a precise attribution of observed metabolic outcomes to the doses and durations of laboratory treatment. However, it did not completely compromise our interpretation to the impacts of the habitual intake, as the variations of the habitual intake levels were restricted by the selection criterion and are comparable to laboratory caffeine intake (300 – 600 mg/day, average 474.1 ± 107.5 mg/day). Second, one might consider the dose administered in our study (150 mg 3 times daily) to be rather high. Earlier studies reported an average caffeine intake per capita varying from 16 to 400 mg/day per capita worldwide (51, 53). However, this number can be underestimated in regular consumers by taking the non-consumers into average (53). Furthermore, while consumption coffee and tea were the primary target of many large-scale country-wise surveys, some other caffeinated products are often missing, such as cola, chocolate, and energy drinks, which could also lead to an underestimation. Thus, the outcomes in our study derived from 450 mg of daily intake may be particularly of interest to the regular consumers, especially those who may consume multiple types of caffeinated products. On the other hand, one might consider that quantifying habitual caffeine levels by self-reports may not be accurate, yet earlier studies have repeatedly reported generally good reliabilities between self-report caffeine intake and salivary concentrations of caffeine and paraxanthine unless the self-report amount is > 600 mg/day (54, 55). In addition, one might consider our sample size to be relatively small despite a within-subject design. Our sample size calculations were indeed based on reported effects sizes of caffeine intake mainly on sleep-wake regulatory indices. Furthermore, we attempted to reduce the variance derived from the sex differences by only including male participants. Thus, the generalizability of the outcomes to female caffeine consumers is limited. Lastly, due to a convenient advantage taken from an existing study, the sampling timing of salivary concentrations may not be optimal, especially a relatively later first sample after the caffeine intake (∼105 minutes). This can potentially bias the calculation of kinetics. It is of importance to note that the kinetics of caffeine are rather comparable to the earlier evidence as discussed previously. Paraxanthine, on the other hand, did not suffer from this issue, as its concentration peaked much later than the earliest sample after the caffeine intake.

### Significance

Caffeine is consumed on a daily basis among 80% of the worldwide population (53). It is of importance to beware that a daily consumption, even merely in daytime, can accumulate the exposure to the psychostimulant and prevents the body from full recovery. The in-progress recovery from a reduced GM and the elevated CBF after 36 h of caffeine cessation entails a longer time required for full restoration than the conventional repetition of daily intake. On the other hand, the accumulation in paraxanthine underscores the importance to investigate its cognitive and physiological effects, which may be responsible for long-term outcomes of chronic exposure to caffeine. Methodologically, the adapted metabolism also suggests a careful consideration to translate acute effects of caffeine onto daily usage. Lastly, the responses of GM and CBF in both the caffeine and withdrawal conditions emphasize the importance of restricting caffeine intake when studying cerebral morphometry and neurovascular activities.

## Acknowledgement

We express our sincere appreciation to our interns Andrea Schumacher and Laura Tincknell, M.Sc. student Sven Leach as well as all the study helpers for their assistance in the experiment and data processing. We also thank Dr. Helen Slawik and Dr. Martin Meyer for the health check during screening process. We especially appreciate all our participants for their volunteering and cooperation.

This manuscript will be published as a preprint on bioRxiv and included in the doctoral thesis of Yu-Shiuan Lin, entitled “*The Influence of Daily Caffeine Consumption on Human Brain Morphometry and Neurocognition*” at the Faculty of Medicine, University of Basel.

## Funding source

Swiss National Science Foundation (grant 320030-163058).

## Conflict of interest

The authors declare no conflict of interest in this study.

## Notes

### Competing Interest Statement

The authors have declared no competing interest.

