## Supplementary material for "Time to recover from daily caffeine intake": Table S1

### Supplement

**Table S1. Mean and standard deviation of concentrations of caffeine (CA) and paraxanthine (PX) in three conditions (μg/ml).**

| | placebo | | caffeine | | withdrawal | |
|---|---|---|---|---|---|---|
| | CA | PX | CA | PX | CA | PX |
| Sample 1 | 0.03 ± 0.03 | 0.11 ± 0.30 | 1.18 ± 1.24 | 5.13 ± 4.00 | 0.46 ± 0.82 | 2.40 ± 3.05 |
| Sample 2 | 0.03 ± 0.04 | 0.09 ± 0.21 | 3.06 ± 1.35 | 6.10 ± 4.11 | 0.42 ± 0.60 | 1.97 ± 2.38 |
| Sample 3 | 0.02 ± 0.01 | 0.07 ± 0.15 | 2.63 ± 1.50 | 5.90 ± 3.65 | 0.34 ± 0.49 | 1.79 ± 2.37 |
| Sample 4 | 0.04 ± 0.09 | 0.07 ± 0.12 | 4.62 ± 2.66 | 7.00 ± 3.81 | 0.25 ± 0.33 | 1.47 ± 1.79 |
| Sample 5 | 0.03 ± 0.05 | 0.06 ± 0.10 | 3.64 ± 1.94 | 8.10 ± 4.98 | 0.20 ± 0.30 | 1.22 ± 1.66 |
| Sample 6 | 0.03 ± 0.04 | 0.05 ± 0.09 | 4.81 ± 2.31 | 8.15 ± 4.22 | 0.17 ± 0.24 | 0.87 ± 1.03 |
| Sample 7 | 0.03 ± 0.04 | 0.06 ± 0.13 | 3.67 ± 2.08 | 8.81 ± 5.08 | 0.12 ± 0.18 | 0.76 ± 0.96 |
| Sample 8 | 0.02 ± 0.02 | 0.05 ± 0.08 | 3.22 ± 2.16 | 8.78 ± 5.03 | 0.10 ± 0.15 | 0.72 ± 1.00 |
| Sample 9 | 0.02 ± 0.01 | 0.04 ± 0.06 | 1.95 ± 1.20 | 7.50 ± 6.79 | 0.06 ± 0.08 | 0.53 ± 0.72 |
| Sample 10 | 0.02 ± 0.01 | 0.03 ± 0.04 | 2.22 ± 2.43 | 7.50 ± 8.98 | 0.06 ± 0.08 | 0.41 ± 0.56 |
| Sample 11 | 0.02 ± 0.00 | 0.03 ± 0.04 | 1.77 ± 1.73 | 6.65 ± 7.90 | 0.05 ± 0.06 | 0.37 ± 0.51 |
